# Selection of trait-specific markers and multi-environment models improve genomic predictive ability in rice

**DOI:** 10.1101/482109

**Authors:** Aditi Bhandari, Jérôme Bartholomé, Tuong-Vi Cao, Nilima Kumari, Julien frouin, Arvind Kumar, Nourollah Ahmadi

## Abstract

Developing high yielding rice varieties that are tolerant to drought stress is crucial for the sustainable livelihood of rice farmers in rainfed rice cropping ecosystems. Genomic selection (GS) promises to be an effective breeding option for these complex traits. We evaluated the effectiveness of two rather new options in the implementation of GS: trait and environment-specific marker selection and the use of multi-environment prediction models. A reference population of 280 rainfed lowland accessions endowed with 215k SNP markers data was phenotyped under a favorable and two managed drought environments. Trait-specific SNP subsets (28k) were selected for each trait under each environment, using results of GWAS performed with the complete genotype dataset. Performances of single-environment and multi-environment genomic prediction models were compared using kernel regression based methods (GBLUP and RKHS) under two cross validation scenario: availability (CV2) or not (CV1) of phenotypic data for the validation set, in one of the environments. The most realistic trait-specific marker selection strategy achieved predictive ability (PA) of genomic prediction was up to 22% higher than markers selected on the bases of neutral linkage disequilibrium (LD). Tolerance to drought stress was up to 32% better predicted by multi-environment models (especially RKHS based models) under CV2 strategy. Under the less favorable CV1 strategy, the multi-environment models achieved similar PA than the single-environment predictions. We also showed that reasonable PA could be obtained with as few as 3,000 SNP markers, even in a population of low LD extent, provided marker selection is based on pairwise LD. The implications of these findings for breeding for drought tolerance are discussed. The most resource sparing option would be accurate phenotyping of the reference population in a favorable environment and under a managed drought, while the candidate population would be phenotyped only under one of those environments.

## Introduction

Drought stress is a major constraint for rice production in rainfed rice growing ecosystems that represent approximately 45% of rice growing areas [1]. It is estimated that 50% of world’s rice production is vulnerable to drought [2]. To reduce yield losses of rice crop in rainfed lowland areas and to increase overall rice production, rice cultivars with improved drought tolerance need to be developed. Early studies reported low selection efficiency for improving grain yield under drought stress [3-4]. Consequently, initial breeding efforts targeted secondary traits such as root architecture, leaf water potential, panicle water potential, osmotic adjustment and relative water content [5-7]. However, these traits do not always have higher broad-sense heritability than grain yield under drought and are not always highly correlated with grain yield [8-9]. Several studies in the last decade have documented the effectiveness of direct selection for grain yield under drought stress [10, 11]. A number of quantitative trait loci (QTL) involved in response of rice grain yield to drought stress have been identified [12] and successfully transferred to elite materiel through marker-assisted selection [13-14]. However, the complex genetic architecture of grain yield under drought hamper the objective of reducing the rice yield loss under drought and achieve higher genetic gain for grain yield under drought.

Genomic selection (GS) has recently emerged as an alternative option to conventional marker-assisted selection targeting mapped QTLs [15-17]. By shifting the plant breeding paradigm from “breeding by design” to a “genome-wide-approach”, GS provides genomic estimated breeding values (GEBV) based on all available marker data, instead of focusing on a limited set of markers that tag putative genes or QTLs. In the last few years, successful proof of GS concept has been reported in maize [18-20], wheat [21-25], barley [26-28] and oats [29]. In rice, moderate to high predictive ability (PA) of GEBV has been reported for a variety of quantitative traits in experiments with the progeny of bi-parental crosses and in diverse germplasm collections [30-35]. In particular, it was shown that rice diversity panels provide accurate genomic predictions for complex traits in the progenies of biparental crosses involving members of the panel [36].

One major feature of all the above-mentioned genomic prediction studies is the use of phenotypic data from only one environment, usually non-stressed. The main reason for such focalization was the lack of appropriate statistical framework that model G×E interactions for the purpose of genomic prediction. A number of such frameworks have recently been proposed. First, the single-trait single-environment genomic best linear unbiased prediction (GBLUP) model was extended to multi-environment context [37]. Then a GBLUP-type model using marker × environment interactions (M×E) was proposed [38]. The M×E based approach was further developed, using a non-linear (Gaussian) kernel and an empirical Bayesian method to model the G×E [39], and extended using Bayesian ridge regression priors that produce shrinkage [40]. The latest models go beyond the extension of single environment models and propose multi-environment prediction models in which the genetic effects are modeled by the Kronecker product of the variance-covariance matrix of genetic correlations between environments with the genomic relationship between lines, using GBLUP or RKGK methods [41]. Application of these multi-environment models to data from multilocal trials of CIMMYT’ maize and wheat breeding programs confirmed their superiority over the single environment models. Multi-environment prediction models were also reported to be better than single environment models when used with rice phenotypic data from managed environments representing two water management systems: continuous flooding and alternate watering and drying [42]. However, the differences between the PA of different categories multi-environment models were not significant. Likewise, it was shown that single-step, best linear unbiased prediction-based reaction norm models using data from non-genotyped and genotyped progenies, can enhance PA in rice recurrent genomic selection [43].

Another feature shared by almost all genomic prediction studies is the use of unsorted markers or markers selected on the base of variety of criteria except association with the target trait. In accordance with the infinitesimal model of the genetic architecture of traits [15], the prediction models are trained and GEBV are computed using the same set of markers for all the phenotypic traits targeted by the breeding programs, whatever their genetic architecture. Zhang *et al.* [44] computed the PA of different genomic prediction methods trained with a relationship matrix built with markers of equal effect (infinitesimal model) and with the same set of markers with weighted effects. The markers’ weight was derived from the result of a ridge regression best linear unbiased prediction or from BayesB prediction for different phenotypic traits. Genomic prediction with markers of weighted effect had higher PA. Similar improvement of the PA of genomic prediction of complex traits was reported, when a trait specific relationship matrix was built using the results of genome wide association studies (GWAS) available in the literature [45].

Here, we report the results of our investigations on the effectiveness of multi-environment prediction models and of trait-specific marker selection in improving the PA of genomic predictions for drought tolerance in rice. We disposed of a large genotypic dataset of 215 k SNP for the reference population (280 accessions) of the rainfed-lowland rice breeding program of the International Rice Research Institute (IRRI), and of phenotypic data of the same population under one normal and two managed drought growth conditions. Subsets of trait and environment-specific markers (28k SNP) were selected for each trait under each the three growth environments, using results of GWAS performed with the complete genotypic dataset. Performances of single-environment and multi-environment genomic prediction models were compared using kernel regression based methods (GBLUP and RKHS).

## Material and methods

Rice populations studied

The plant material was a diversity panel of 280 accessions, of which 245 belong to the *indica* genetic group and 35 to the *aus* group (S1 Table). Landraces originating from South and South-East Asia represented the largest share of the panel (215). The remaining 65 accessions were improved lines. The 280 accessions were selected for their potential interest for the IRRI rainfed lowland rice breeding program, among the 3,000 rice accessions recently re-sequenced [46]. Seeds of the plant material for the present study were provided by the IRRI Gene-bank.

Phenotyping and analysis of phenotypic data

### Field experiments

Three independent experiments were conducted in the 2015 dry season at IRRI (14.18°N, 121.25°E), Philippines, to evaluate the performance of the 280 accessions under three managed environments, E1, E2 and E3. E1 corresponded to the standard lowland rice cultivation, without stress, i.e. transplanting crop establishment in puddled soil and continuous flooding (anaerobic conditions) throughout the crop cycle. E2 corresponded to standard lowland rice cultivation associated with application of drought stress at the reproductive stage (see below). E3 corresponded to standard cultivation of upland rice (crop establishment by direct seeding in well-drained soil with continuous aerobic conditions during the crop cycle) with drought stress applied at the reproductive stage.

Drought stress at the reproductive stage was applied following the procedure described in [13]. Briefly, in the E2 experiment, the field was drained 30 days after transplanting and irrigation was withheld until severe leaf rolling was observed in at least 75% of the accessions. Fields were thereafter re-irrigated by sprinklers and the water was again drained after 24 hours in another cycle of drought stress. This cyclic pattern was continued until harvest. In the E3 experiment, drought stress was started 45 days after sowing, by withholding sprinkler irrigation until the soil water tension fell below –50 kPa at a depth of 30 cm. Thereafter, drought stress was applied in a cycle of sprinkler-irrigation and drainage 24 hours later until harvest.

The experimental design in E1 was an augmented randomized complete block, with 16 checks, in single row plots with rows 5 m in length. The E2 and E3 experiments were planted in an Alpha-lattice design with 2 replications and 16 checks, in two-row plots with rows 5 m and 3 m in length, respectively.

Three traits were measured in or computed for each individual plot. Days to flowering (DTF, day) as the number days between sowing and visual estimation of 50% exerted panicles of the plot.Plant height at maturity (PH, cm), was measured as the mean height from the soil surface to the tip of the panicle of the main tiller of three randomly chosen plants. Grain yield (GY, kg/ha), was computed from the grain weight at 12.5% moisture of five plants harvested at physiological maturity.

### Analysis of phenotypic data

The drought stress applied in E2 and E3 experiments proved to be too sever for a number of accessions that did not reach the reproductive stage or did not produce any grain. The three traits could consequently only be measured properly in 230 accessions in E2 and in 226 accessions in E3, of which 204 were common to the two experiments. Consequently, data analyses were run for 280 and 204 accessions in E1 and for 204 accessions in the E2 and E3.

The phenotypic data were analyzed using the PROC MIXED procedure of SAS v9.0 (SAS Institute Inc., Cary NC, USA).

For each experiment and trait, the following model was used to adjust phenotypic measurements:

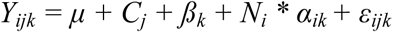

where *Y*_*ijk*_ is the phenotype of either the *i*^*th*^ accession or the *j*^*th*^ check in the *k*^*th*^ block, *μ* the overall mean, *C*_*j*_ the checks effect considered as fixed, *μ*_*k*_ the block effect considered as random, *N*_*i*_ a covariate that amounted one for accessions and zero for checks entrees in each block, *α*_*ik*_ the random effect of the *i*^*th*^ accession in the *k*^*th*^ block and *ε*_*ijk*_ is the residual considered as random. The block effects were estimated using data from the 16 checks.

The variance component was estimated using the restricted maximum likelihood method. Finally, Y_adj_ values were extracted for each genotype and were used in genomic prediction models.

For each trait, broad sense heritability was calculated using the ratio H^2^ = s^2^ _g/s ^2^ p_, where s^2^ ^g^ is the genotypic variance obtained from the experimental data and the phenotypic variance s^2^ p= s^2^ g + s^2^ e, where s^2^ eis the residual variance obtained from the model.

## Genotyping and analysis of genotypic data

### Genotypic data

The genotypic data for the 280 accessions were obtained from the International Rice Informatics Consortium (IRIC) database for the 3,000 rice genomes project (http://iric.irri.org/). It corresponded to the 962k Core-SNP dataset. Filtering of the 962k loci for missing data (threshold of ≤ 20%), the frequency of the minor allele (MAF), threshold of ≥ 2%, and for the rate of heterozygosity (Ho, threshold of ≤ 5%), resulted in a working set of 215,242 SNP. The average rate of missing data per locus for the 280 accessions was 4.1%. The rate of missing data per accession ranged from 1.5 to 17.3%. Heterozygoty (Ho) per accession was 0.2 to 10.5, average 1.0%. The average minor allele frequency was 14.6%. The missing data were imputed using Beagle v4.0 [47]. The final matrix of 280 accessions and 215,242 SNP is hereafter referred to as the “215k SNP” marker set.

### Characterization of population structure and linkage disequilibrium

The genetic diversity of the panel was investigated using 4,824 SNP with no missing data. First pairwise dissymmetry between genotypes was computed using simple matching information and an unweighted neighbor-joining tree was built using Darwin software v6 [48]. Then the tree topology was visualized using FigTree v1.4.3. [49]. Pairwise linkage disequilibrium (LD) between SNP loci was calculated at the level of the individual chromosome, using the working genotypic datasets and the r^2^ estimator proposed for non-phased genotypic data [50].

### Marker selection for genomic prediction

Taking advantage of the large quantity of genotypic data available, three marker-selection approaches were tested, with the aim of (i) identifying the minimum marker density while maintaining the highest possible PA of gnomic predictions and (ii) of analyzing the performances of trait-specific marker selection.

In the first approach, five LD thresholds (r^2^ ≤ 0.25, r^2^ ≤ 0.50, r^2^ ≤ 0.75, r^2^ ≤ 0.90, and r^2^ ≤ 1) combined with three MAF thresholds (≥ 2%, ≥ 5% and ≥ 25%) were considered based on the following procedure. The complete pairwise r^2^ matrix for each chromosome was computed on the 215k SNP dataset. Then for each chromosome, single loci or clusters of loci with pairwise LD with other loci below the threshold were identified. All singletons (LD with other loci below the given threshold) were kept. For the clusters of loci, one locus with the fewest missing data before imputation and the highest MAF was randomly selected to represent the cluster. Lastly, the three threshold of MAF were applied to each of the five levels of LD, leading to 15 incidence matrices, the smallest composed of 2,859 SNP (S2 Table). Comparison of PA of genomic predictions obtained with each of these matrices, using phenotypic data from E1 environment, (see below and results section) identified the matrix with 28,091 SNP (r^2^ ≤ 0.5 and MAF ≥ 5%), hereafter referred to as 28k SNP dataset, as the best compromise.

In the second approach, the selection criterion was the degree of association of the SNP loci with the target phenotypic trait, as measured by genome-wide association analysis (GWAS). Taking the results of the first selection into account, marker selection was performed as follow: (i) the 215k SNP dataset was filtered for MAF ≥ 5%. (ii) The resulting set of 148,916 SNP was used for GWAS, for each trait, under each of the three environments (E1, E2 and E3), using the standard mixed linear model (MLM) under Tassel v5.2.48 (Bradbury et al; 2007). MLM uses a genotype based kinship matrix (K) jointly with population structure (Q). We computed K and Q (coordinate of the accessions on the five first axis of a principal component analysis) using the 148,916 SNP data set. (iii) For each trait, under each drought management, 28,091 SNP with the smallest P-value of trait-marker association were extracted to serve in genomic prediction experiments.

The third approach differed from the second in that the GWAS experiments were implemented with accessions that were not included in the corresponding model training process for the genomic prediction experiments. The objective was to avoid overfitting of the training model. The practical implication was the need to select a new set of 28k SNP for each of the replicates of each genomic prediction experiment (see below).

Hereafter, we refer to the first marker selection approach as LD-based selection, to the second approach as GWAS-based selection and to the third approach as S-GWAS-based selection.

## Statistical methods for genomic prediction

### Single environment models

Two kernel regression models were used to predict GEBV in each drought stress experiment: the genomic best linear unbiased prediction (GBLUP) and the reproducing kernel Hilbert spaces regressions, RKHS, [51]. The implementation procedures of the two models is detailed in [42]. Briefly the GBLUP method that hypothesizes a strictly additive determinism of the genetic effects [52] was implemented using the genomic matrix G = M*M’ (M being the incidence matrix) and the Expectation-Maximization convergence algorithm with the R package *KRMM* [53]. The RKHS method that captures more complex genetic determinism [54] was also implemented using the *KRMM*.

### Multi-environment models

To predict the GEBV with data from the three environments (E1, E2 and E3), hereafter referred to as multi-environment prediction, we used the statistical models described above with extensions to include environmental effects. In the extended GBLUP model, the effects of environments and markers are separated into two components: the main effect of the markers for all the environments and the effect of markers for each individual environment [38]. For RKHS, we used the extended models incorporating G×E corresponding to the “Empirical Bayesian–Genotype **×** Environment Interaction Model” proposed in [39].

### Implementation of the models

Analyses were performed in the R 3.4.2 environment [55] with the R packages *BGLR* 1.0.5 (Pérez and de los Campos 2014) and *MTM* 1.0.0 (De los Campos and Grüneberg 2018). For both packages, 25,000 iterations for the Gibbs sampler were used after 5,000 burn-ins were removed. The analyses were supported by the CIRAD - UMR AGAP HPC Data Center of the South Green Bioinformatics platform (http://www.southgreen.fr/).

### Assessing predictive ability of genomic prediction

In the single environment model, 80% of the accessions (i.e. 224 and 163 for the population of 280 and 204 accessions, respectively) were used as the training set and the remaining 20% (56 and 41, for the population of 280 and 204 accessions, respectively) was used as the validation set.

The multi-environment models were implemented for the three possible 1×1 combinations of the three environments [E1(E2), E1(E3), E2(E1), E2(E3), E3(E1), and E3(E2)] and the three possible 2×1 combinations, [E1(E2 + E3), E2(E1 + E3) and E3(E1 + E2)]. In each case, the PA was assessed with two different cross-validation schemes. The first method (CV1), which resembled what was done in the single environment, used 80% of the observations as a training set and the remaining 20% as the validation set and assumed that phenotypic observations for the two or three environments are available for the individuals serving as training set, and that no phenotypic data are available for the individuals in the validation set. CV1 corresponds to the situation in which the phenotypes of newly generated individuals have to be predicted based only on their genotypic information [38]. The second method (CV2) also used 80% of the observations as a training set and the remaining 20% as the validation set but assumed that at least one observation in one environment was available for the individuals in both the training set and the validation set [38]. The same proportions of the training set and validation set (i.e. 80% and 20%, respectively) were kept when the prediction model combined data from the three environments.

Under both single and multi-environment models, one hundred replicates were computed for all random partitioning in the training and validation sets. The PA of each partition was calculated as the Pearson correlation coefficient between predictions and phenotypes in the validation set. The resulting estimates of PA were averaged and the associated standard error was calculated. For the multi-environment models, the correlation was calculated within each environment. For each trait (DTF, GY and PH) and each statistical method (GBLUP, RKHS), the same partitions were used to compute the PA. The number of replicates under S-GWAS marker selection method was limited to 20.

The correlation coefficient data of all prediction experiments were transformed into a Z-statistic using the equation: Z = 0.5{ln[1 + r]−ln[1 − r]} and analyzed as a dependent variable in an analysis of variance. After estimation of confidence limits and means for Z, these were transformed back to r variable. In each case, ANOVA was performed to partition the variance of PA into different sources, with all effects declared as fixed.

## Results

### Phenotypic characteristics of the populations

Box-plots of the adjusted means of the three phenotypic traits under the three environments are presented in Fig 1. As expected, average DTF increased under the drought environment E2 and E3. Conversely, GY and PH decreased. Drought also affected phenotypic variances, which decreased for GY and PH, and increased for DTF. However, in all case, the distribution of each trait in each environment was reasonably symmetric.

**Fig 1.**
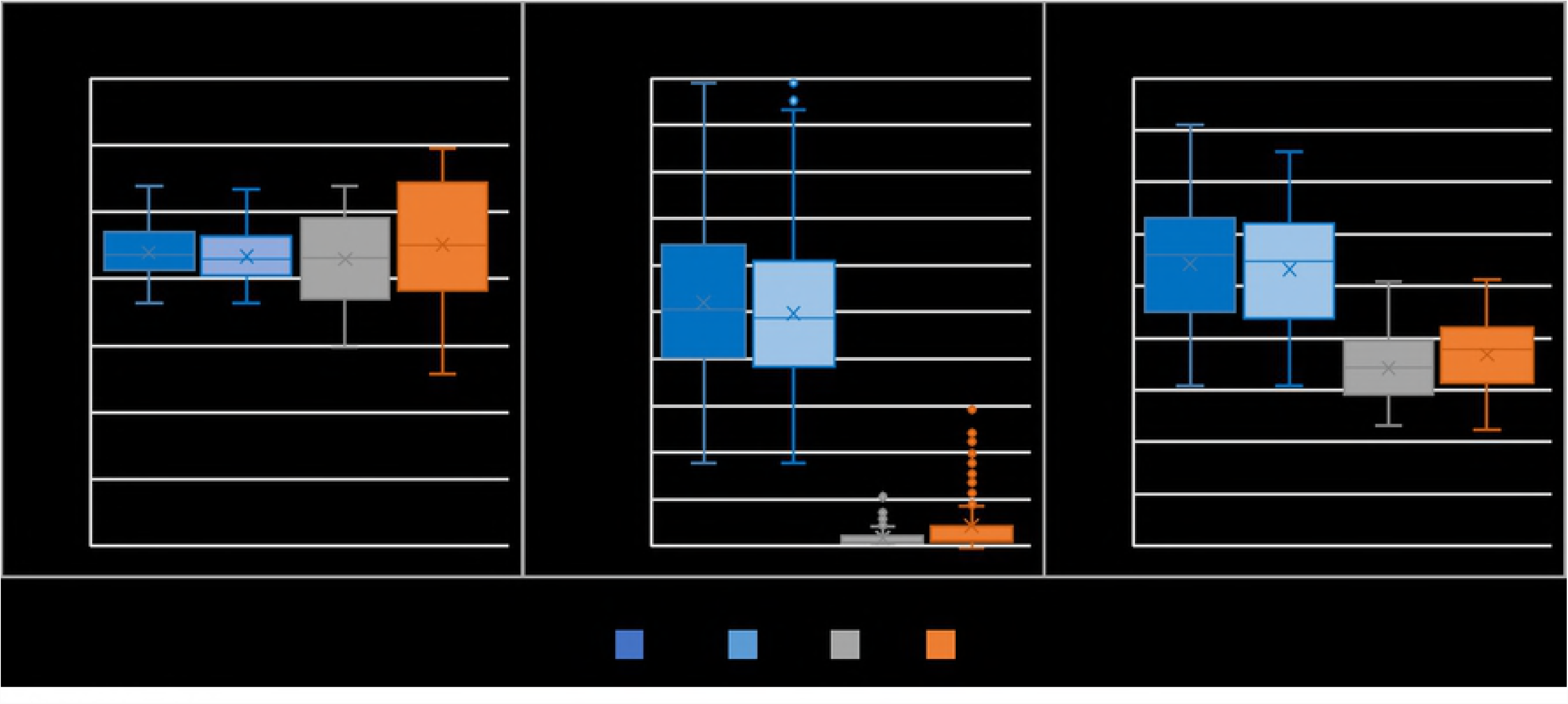
Boxplot of phenotypic variables within the rice population. E1c and E1: data from lowland-non-stressed stressed environment with 280 and 204 accessions, respectively. E2 and E3: data from lowland-drought and upland-drought environments, respectively, with 204 accessions.

Correlation between the adjusted means under non-drought (E1) and drought environments (E2 and E3) was above 0.50 and highly significant for DTF and PH, close to zero for GY (Table 1). Correlations between the adjusted means under E2 and E3 ranged between 0.50 for GY and 0.74 for PH. Overall, the pattern of these correlations suggests significant G×E interactions for response to drought (E2 and E3 compared to E1), much less G×E interactions between lowland and upland drought (E2 compared to E3).

**Table 1:** Analysis of factors that influence the variation in predictive ability of single and multi-environment prediction models. The effects of the statistical model (GBLUP, RKHS-1 and RKHS-2), the trait (DTF, GY and PH), the cross-validation strategy (CV1 and CV2), drought stress environments (E1, E2 and E3) and different combinations of training environment and their interactions were evaluated.

| **Analysis with only main effects** |  |  |  |  |  |
| --- | --- | --- | --- | --- | --- |
| **Source** | **DF** | **SS** | **MS** | **F-value** | **ProbF** |
| Model | 10 | 309.728 | 30.973 | 1 773.667 | < 0.0001 |
| Error | 32 389 | 565.595 | 0.017 |  |  |
| Corrected Total | 32 399 | 875.323 |  |  |  |
| Trait | 2 | 145.581 | 72.791 | 7 141.750 | <.0001 |
| Model | 1 | 10.730 | 10.730 | 1 052.770 | <.0001 |
| Method | 2 | 1.426 | 0.713 | 69.930 | <.0001 |
| CV | 1 | 15.993 | 15.993 | 1 569.160 | <.0001 |
| Drought environment | 2 | 134.409 | 67.204 | 6 593.650 | <.0001 |
| Training environment | 2 | 1.588 | 0.794 | 77.880 | <.0001 |
| **Analysis with main effects and all first-order interactions** |  |  |  |  |  |
| Model | 51 | 496.946 | 9.744 | 833.032 | < 0.0001 |
| Error | 32 348 | 378.377 | 0.012 |  |  |
| Corrected Total | 32 399 | 875.323 |  |  |  |
| Trait | 2 | 145.581 | 72.791 | 7 141.750 | <.0001 |
| Model | 1 | 10.730 | 10.730 | 1 052.770 | <.0001 |
| Method | 2 | 1.426 | 0.713 | 69.930 | <.0001 |
| CV | 1 | 15.993 | 15.993 | 1 569.160 | <.0001 |
| Drought environment | 2 | 134.409 | 67.204 | 6 593.650 | <.0001 |
| Training environment | 2 | 1.588 | 0.794 | 77.880 | <.0001 |
| Trait*Model | 2 | 4.605 | 2.303 | 225.920 | <.0001 |
| Trait*Method | 4 | 1.418 | 0.355 | 34.780 | <.0001 |
| Model*Method | 2 | 1.812 | 0.906 | 88.910 | <.0001 |
| Trait*CV | 2 | 1.882 | 0.941 | 92.330 | <.0001 |
| Model*CV | 1 | 15.993 | 15.993 | 1 569.160 | <.0001 |
| Method*CV | 2 | 1.627 | 0.814 | 79.820 | <.0001 |
| Trait*Drought | 4 | 152.543 | 38.136 | 3 741.620 | <.0001 |
| Model*Drought | 2 | 0.786 | 0.393 | 38.580 | <.0001 |
| Method*Drought | 4 | 0.499 | 0.125 | 12.240 | <.0001 |
| CV*Drought | 2 | 0.792 | 0.396 | 38.840 | <.0001 |
| Trait* Training environment | 4 | 0.957 | 0.239 | 23.470 | <.0001 |
| Model* Training environment | 2 | 1.588 | 0.794 | 77.880 | <.0001 |
| Method* Training environment | 4 | 0.935 | 0.234 | 22.930 | <.0001 |
| CV* Training environment | 2 | 1.782 | 0.891 | 87.440 | <.0001 |
| Drought* Training environment | 4 | 1.686 | 0.422 | 41.350 | <.0001 |

DF: Degree of freedom; SS: Sum of squares; MS: Mean square.

Partitioning of the observed phenotypic variation into different sources of variation via the mixed model analysis is shown in **Error! Reference source not found.**. Whatever the trait or drought environment, the genotype contributed significantly to the phenotypic variation. The highest contribution of the genotype effect to the phenotypic variation, relative to residual, was observed for PH under E1 and for DTF under E3. Whatever the environment, the genotype effect contributed the least to GY variations. The broad-sense heritability *H*^*2*^ was rather high even for GY. *H*^*2*^ ranged from 0.75 to 0.86 for DTF, from 0.68 to 0.91 for GY, and from 0.81 to 0.96 for PH. Given the differences in the experimental design in the three environments, combined analysis of data was performed using accessions’ adjusted mean from each environment. The genotype effect was highly significant for the three traits. This was also the case for the residual variance that included the phenotypic variance due to G×E interactions.

### Genotypic characteristics of the population

Within the 215k genotypic dataset, the average marker density along the 12 chromosomes was one marker every 1.95 kb. However, the SNP loci were unevenly distributed (S1 Fig). The 28k genotypic dataset represented an average marker density of one marker every 13.7 kb. However, 426 pairs of adjacent loci had distances above 100 kb, the largest distance being 427 kb.

The distribution of MAF was skewed toward low frequencies in both 215k and 28k datasets. The MAF was below 10% for 50% and 37% of loci in 215k and 28k datasets, respectively.

The average LD was rather low, 0.123 and 0.063 in the 215k and 28k datasets, respectively. The decay of LD along physical distance is presented in the (S2 Fig). For between-marker distances of 0 to 25 kb, the average r^2^ value was 0.145 and 0.103 in the 215k and 28k datasets, respectively. These r^2^ dropped to half their initial level at a mean distance between markers of around 600 kb and 250 kb, in the 215k and 28k datasets, respectively.

Analysis of genetic diversity confirmed the initial assignment (see Materiel and Methods) of accessions to *indica* and *aus* groups as well as the presence of subgroups within the *indica* group (S3 Fig).

### Effect of characteristics of the incidence matrix on predictive ability of genomic prediction

The effect of the characteristics of the incidence matrix was analyzed using phenotypic data of the reference population of 280 accessions, under the E1 environment. The 90 cross validation experiments involving five levels of LD, three levels of MAF, two prediction methods and three phenotypic traits yielded mean PA of genomic prediction ranging from 0.417 to 0.586 (Fig 2; S3 Table A). Among these sources of variation, only the effects of phenotypic trait and LD were significant as well as the interaction between those factors (S3 Table B). The highest average PA was observed for DTF (r = 0.566) and the lowest for GY (r = 0.427). The Fisher least significant difference (F-LSD) indicated that, whatever the trait, the incidence matrices with LD value of r^2^ ≤ 0.25 led to significantly higher PA than the incidence matrices with r^2^ > 0.25, except for DTF. In the latter case, the difference in PA between r^2^ ≤ 0.25 and r^2^ ≤ 0.5 was not significant (S3 Table B). Given these results, and the fact that the next steps of our study dealt with phenotypic data from three environments, we took the conservative decision of using the 28k SNP dataset (r^2^ ≤ 0.50, MAF ≥ 5%) as the reference size for the incidence matrices.

**Fig 2.**
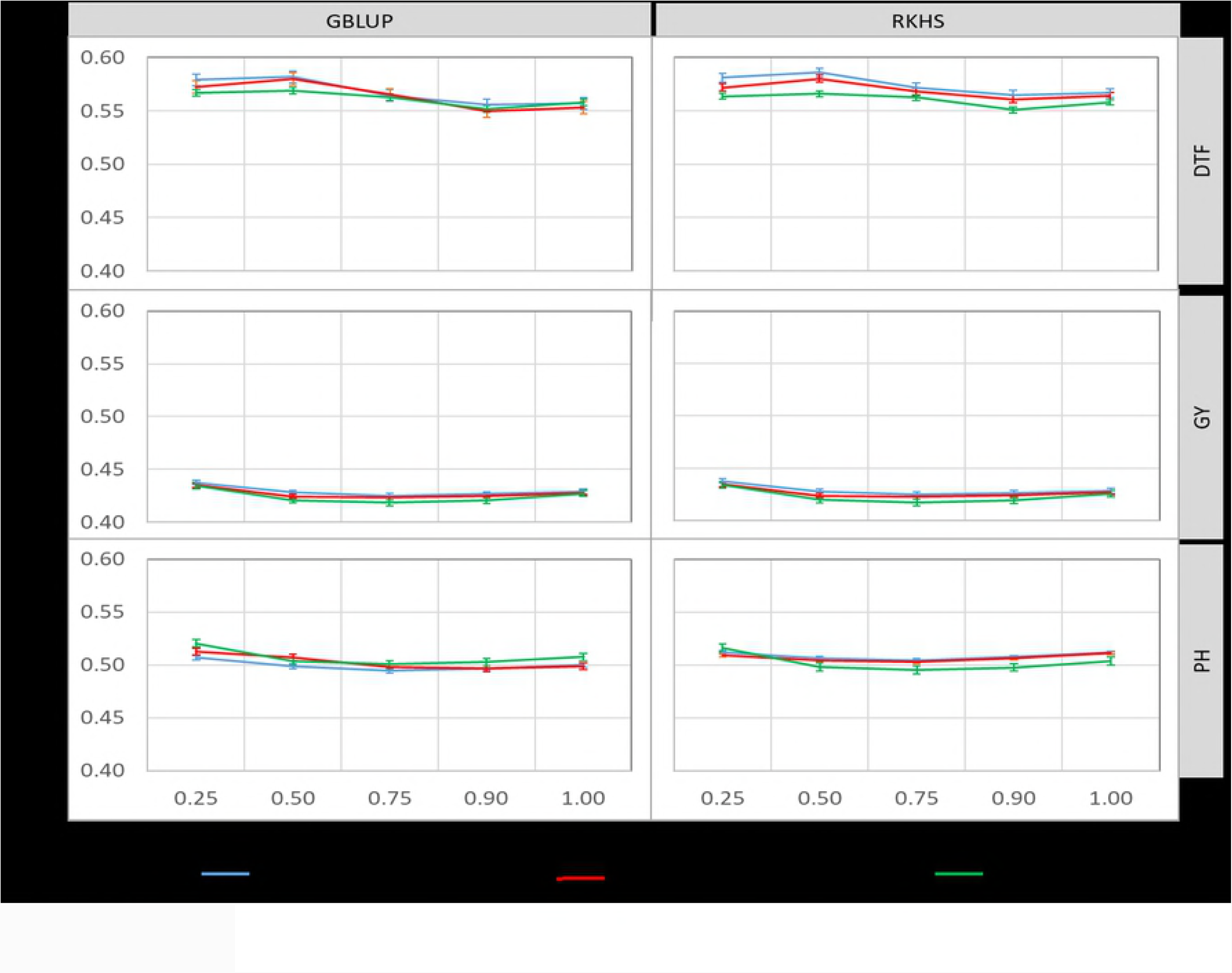
Predictive ability of genomic prediction for combinations of five levels of linkage disequilibrium (LD) and three levels of minor allele frequency (MAF) in cross validation experiments in the rice population of 280 accessions, under non-stress condition (E1), for days to flowering (DTF), grain yield (GY) and plant height (PH), obtained with 2 statistical methods, GBLUP and RKHS.

### Effect of trait and environment-specific marker selection on predictive ability of genomic prediction

The effect of trait specific selection of the 28k markers on the predictive ability of genomic prediction was analyzed in the reference population of 280 accessions under the E1 environment. Both GWAS-derived and S-GWAS-derived markers led to significantly higher PA than LD-derived markers for the three traits considered (Table 2). Under GWAS-derived markers, the effects of trait, the prediction method and their interactions were also significant (S4 Table). Compared to LD-derived markers, the GWAS-derived markers led to average PA gain of 0.55% for DTF, 111% for GY and 67% for PH. The average PA gains were lower with S-GWAS-derived markers: 16% for DTF, 26% for GY and 24% for PH.

**Table 2:** Predictive ability (AP) of GBLUP and RKHS statistical models obtained with 28,091 SNPs selected on the basis of linkage disequilibrium (LD), of genome wide association study results (GWAS) and in astringent GWAS (S-GWAS) results.

|  |  |  | Environment & size of the population |  |  |  |  |  |  |  |  |  |  |  |
| --- | --- | --- | --- | --- | --- | --- | --- | --- | --- | --- | --- | --- | --- | --- |
|  |  |  | 280 accessions |  |  |  |  |  | 204 accessions |  |  |  |  |  |
|  |  |  | E1 |  |  | E1 |  |  | E2 |  |  | E3 |  |  |
|  |  |  | LD | GWAS | S-GWAS | LD | GWAS | S-GWAS | LD | GWAS | S-GWAS | LD | GWAS | S-GWAS |
| GBLUP | DTF | PA | **0.540 b** | **0.850 a** | **0.618 a** | **0.599 b** | **0.892 a** | **0.654 a** | **0.450 b** | **0.513 a** | **0.516 a** | **0.579 b** | **0.713 a** | **0.632 a** |
|  |  | STD | 0.084 | 0.038 | 0.068 | 0.082 | 0.033 | 0.072 | 0.129 | 0.111 | 0.109 | 0.098 | 0.077 | 0.098 |
|  | GY | PA | **0.389 b** | **0.839 a** | **0.489 a** | **0.345 b** | **0.400 a** | **0.396 b** | **0.420 b** | **0.913 a** | **0.478 b** | **0.672 b** | **0.891 a** | **0.684 a** |
|  |  | STD | 0.090 | 0.038 | 0.062 | 0.115 | 0.103 | 0.125 | 0.120 | 0.034 | 0.160 | 0.122 | 0.065 | 0.132 |
|  | PH | PA | **0.505 b** | **0.858 a** | **0.623 a** | **0.558 b** | **0.843 a** | **0.597 a** | **0.648 b** | **0.916 a** | **0.678 b** | **0.673 b** | **0.887 a** | **0.701 a** |
|  |  | STD | 0.087 | 0.038 | 0.086 | 0.085 | 0.054 | 0.045 | 0.087 | 0.025 | 0.049 | 0.073 | 0.041 | 0.058 |
| RKHS | DTF | PA | **0.539 b** | **0.819 a** | **0.633 a** | **0.585 b** | **0.862 a** | **0.660 a** | **0.466 b** | **0.520a** | **0.552 a** | **0.554 b** | **0.697 a** | **0.630 a** |
|  |  | STD | 0.081 | 0.045 | 0.063 | 0.090 | 0.042 | 0.072 | 0.114 | 0.107 | 0.111 | 0.098 | 0.079 | 0.089 |
|  | GY | PA | **0.387 b** | **0.795 a** | **0.490 a** | **0.352 b** | **0.385 a** | **0.392 b** | **0.435 b** | **0.850 a** | **0.481 b** | **0.668 b** | **0.839 a** | **0.693 b** |
|  |  | STD | 0.089 | 0.045 | 0.056 | 0.106 | 0.103 | 0.128 | 0.113 | 0.046 | 0.681 | 0.149 | 0.092 | 0.133 |
|  | PH | PA | **0.501 b** | **0.818 a** | **0.625 a** | **0.547 b** | **0.811 a** | **0.597a** | **0.646 b** | **0.887 a** | **0.681 a** | **0.675 b** | **0.862 a** | **0.712 a** |
|  |  | STD | 0.090 | 0.045 | 0.089 | 0.084 | 0.060 | 0.045 | 0.078 | 0.031 | 0.052 | 0.075 | 0.047 | 0.059 |

DTF: days to flowering; GY: grain yield PH: plant height (PH); E1: lowland non-stress environment; E2: lowland stressed; E3: upland stressed.
AP of GWAS-derived markers and S-GWAS-derived markers based prediction that are followed by different letters than their LD-derived counterpart are significantly different at $\alpha < 0.05$ .

The effect of trait and environment specific selection of the 28k markers on the predictive ability of genomic prediction was analyzed within the population of 204 accessions phenotyped under three drought environments, E1, E2 and E3. Whatever the phenotypic trait, the environment and the genomic prediction method, GWAS-derived markers resulted in systematically significantly higher PA than their LD-derived counterpart (Fig 3, Table 2). The effect of interactions between the marker selection method, the prediction method and environment were also often highly significant (S5 Table). For instance, for DTF under E1, the effect of the marker selection method was significant with the RKHS prediction method but not with GBLUP. The reverse was observed for GY and the interaction of prediction method with environment was not significant for PH. Under E1, the average PA gains with GWAS-derived markers, compared to LD derived markers, were of 29% for DTF, 49% for GY and 40% for PH. These PA gains were much smaller than the ones observed under the some environment with the population of 280 accessions, suggesting an effect of population size. Under E2 and E3, the average gains in PA for the three traits were 53% and 28%, respectively. These PA gains were affected by trait × environment interaction. This was specially the case for GY with PA gain of 13% under E1, 106% under E2 and 29% under E3.

**Fig 3.**
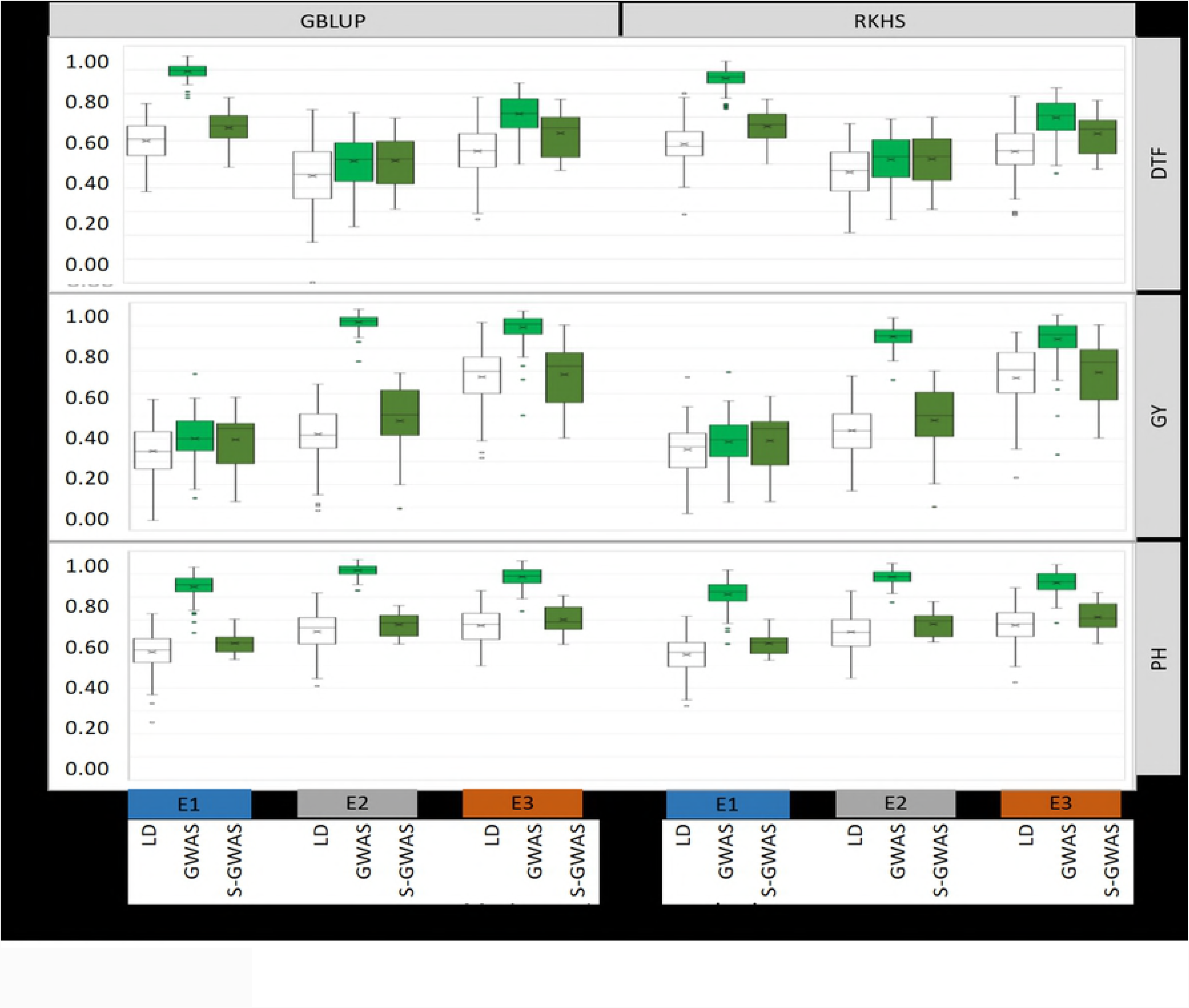
Predictive ability of genomic prediction in cross validation experiments implemented with 28,091 SNP derived using three marker selection methods: Linkage disequilibrium (LD) between markers, white boxes; Genome wide association analysis with the target traits (GWAS), pale green boxes; and astringent genome wide association analysis with the target traits (S-GWAS), dark green boxes. The three traits, days to flowering (DTF), grain yield (GY) and plant height (PH), were phenotyped under three environments: rainfed lowland (E1), rainfed lowland with drought stress (E2) and upland with drought stress (E3). For each box, the mean (x) and median (horizontal bar) values are represented.

The S-GAWS-derived markers also resulted in systematically higher PA of genomic prediction than the corresponding LD-derived markers (Table 2). The overall effect of the marker selection method was systematically significant for each of the three traits considered. Significant interactions between the marker selection method and the environment were observed only for the DTF trait (S6 Table). The average gain in PA, over LD derived markers was of 11% under E1, 10% under E2 and 6% under E3 (Fig 3 and Table 2). Similar to GWAS-derived markers, these gains in PA were much smaller than the ones observed for S-GWAS derived markers in the population of 280 accessions, and were affected by trait × environment interactions. The F-LSD values indicate that the gains of PA for DTF (12% in average) were significant under each of the three environments. For PH, the PA gains (9% in average) were significant only under E1 and E3. For GY, the PA gains (6% on average) were not significant under any of the three environments. It is noteworthy that, whatever the marker selection method, the standard deviation of genomic predictions for GY in the population of 204 accessions, was almost systematically higher (average STD = 0.111) than the ones observed for DTF (average STD = 0.090) and PH (average STD = 0.058).

Compared to GWAS-derived markers, S-GWAS derived markers led to an average PA loss of 15% for DTF, 26% for GY and 24% for PH (Table 2). However, the individual PA differences were strongly affected by the trait × environment interaction. For instance, under the E2 environment, S-GWAS-derived markers performed as well as GWAS-derived markers for DTF while a PA loss of 42% was observed for GY. Conversely, under E1, S-GWAS-derived markers performed as well as GWAS-derived markers for GY while a PA loss of 26% was observed for DTF.

The two prediction methods GBLUP and RKHS (overall average PA of 0.638 and 0.628, respectively, performed similarly. Whatever the environment and the size of the population, the differences in PA between the two methods did not exceed 6%. Likewise, the average difference in PA between the two methods was only less than 1% for DTF and PH, 4% for GY, to the advantage of GBLUP.

### Predictive ability of multi environment genomic prediction

Comparisons of the PA of the multi-environment genomic predictions for the three traits under the two cross validations strategies, three drought environments and two prediction methods are summarized in Fig 4. PA ranged from 0.226 to 0.809. Analysis of the sources of variation of PA revealed a significant effect of all the factors considered, ranked in decreasing importance: the cross-validation strategy, the trait, the environment, the type of model, and the statistical method. Interactions between the factors were also significant (Table 3). Given these interactions, for each trait, each multi-environment prediction was compared to its single environment counterpart, using Student’s test (S7 Table).

**Fig 4.**
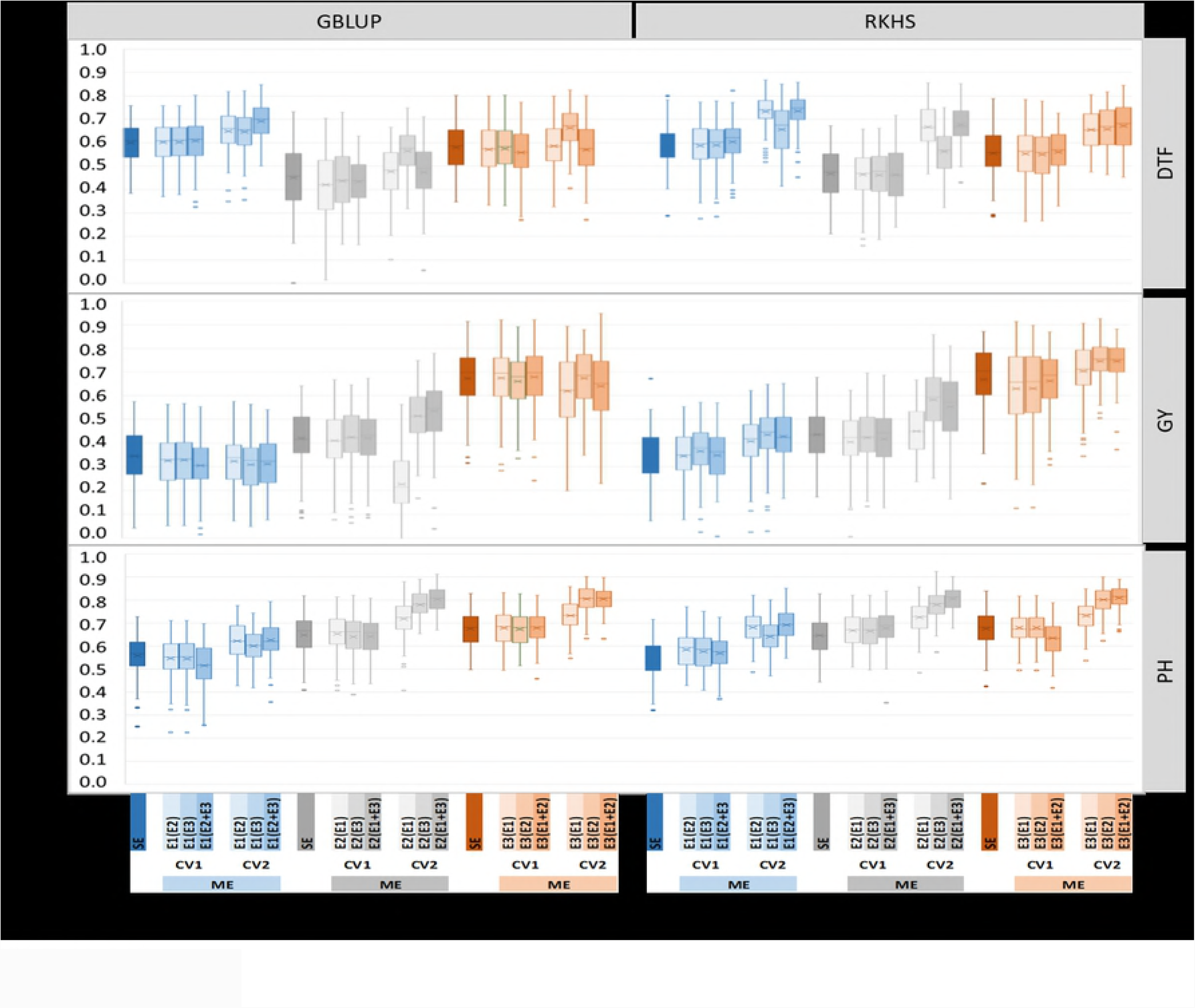
Predictive ability of genomic prediction experiment with single environment (SE), and multi-environment (ME) models obtained with the GBLUP, RKHS-1 and RKHS-2 statistical methods. Traits studied are days to flowering (DTF), grain yield (GY) and plant height (PH). The ME models are implemented with two cross-validation strategies CV1 and CV2. Three environments are considered: lowland no-drought (E1, blue), lowland drought (E2, gray) and upland drought (E3, orange). Environment in brackets contributed to the training of ME models. For each box, the mean (x) and median (horizontal bar) values are represented

Whatever the trait or the environment, the multi-environment models combined with the CV2 strategy provided the highest average PA of 0.630, and outperformed single environment models, with an average gain of 22% for DTF, of 6% for GY and 15% for PH. Under CV2, the multi-environment models using interaction between the non-stressed environment E1 and one of the drought environments E2 or E3, to predict phenotypic performance in E2 or E3 (i.e. E2(E1) or E3(E1)), resulted in similar PA to single environment models. Conversely, the multi-environment model E2(E3) and E3(E2) resulted in PA gains of 22% and 13% respectively compared to single environment model. The multi-environment models predicting phenotypic performances in one of the drought environment, using data from the non-stressed environment and the remaining drought environment (i.e E2(E1 + E3) and E3(E1 + E2)), resulted in PA gains of 24% over the corresponding single environment model.

Performances of the multi-environment prediction models combined with the CV1 strategy (average PA of 0.408) did not significantly differ from their single environment model counterpart, whatever the trait and the environment and the prediction method considered. Multi-environment models predicting performances in drought stressed environments from E2(E1 + E3) and E3(E1 + E2) or from E2(E1) and E3(E1) did not result in significantly different PA either.

Among the multi-environment prediction models, RKHS, with average PA of 0.600 performed systematically slightly better than the gGBLUP, average PA of 0.566.

## Discussion

The drought stress applied in this study was so severe that 25% of the accessions did not reach the flowering stage. However, the stress corresponded to the drought severity rice crops often facing in the drought-prone rainfed lowland ecosystem of South Asia [56]. Despite the severity of the drought stress, the broad-sense heritability observed for the three phenotypic traits was reasonably high, and the distribution of the three phenotypic traits in each of the three environments was reasonability symmetric, once the accessions that did not reach the flowering stage were discarded.

### Effect of the characteristics of the incidence matrix on the predictive ability of genomic prediction

The promise of GS to improve response to selection in populations that are under artificial selection is often presented as depending on the availability of high-density genotypic data to maximize the number of QTLs in LD with at least one marker [15-17], and a large training population to accurately estimate marker effects. However, high-density SNP genotyping may be too expensive for many plant breeding programs, especially when it comes to genotyping of the selection candidates in each breeding cycle. Regarding the size of the population, for a given amount of resources, breeders can choose between evaluating more individuals less accurately or fewer individuals more accurately. Simulation studies have shown the feasibility of using low-density genotypic data, with SNP markers evenly distributed along the genome with no significant decrease in the PA of genomic predictions [57-58]. Empirical studies of PA of genomic prediction in various plant species also showed very little, if any, effect of marker densities, until very low marker densities were used [59]. The results of our analysis of the effects of LD and MAF and the associated marker density confirmed these findings. The highest PA were observed with marker densities below one SNP every 15kb. On the other hand, our results somewhat deviate from the rule of positive relationship between the PA of GEBV and the size of the training set [59-61]. Indeed, we did not observe any significant differences in PA between the populations of 280 and 204 accessions evaluated under the E1 environment. The reasons for this deviation may be that (i) the rule applies mainly in across population predictions whereas we were performing cross-validation within each population; (ii) the reduction in size from 280 to 204 accessions was accompanied by a change in the structure of the populations. Indeed, as all the discarded accessions belonged to the more drought susceptible *indica* group, the proportion of *aus* accessions increased in the population of 204 accessions from 12% to 17%.

Compared to LD based marker selection, GWAS-based and S-GWAS based marker selection led, on average, to much higher PA in cross validation experiments within the population of 280 accessions: 77% and 22%, respectively. The gains in PA were smaller but still statistically significant, in cross validation experiments within the population of 204 accessions: 39% and 6%, respectively. Similar results were reported in a simulation based comparison of GEBV obtained with the GGBLUP method using relationship matrices derived from trait-specific markers or from non-trait-specific markers [43]. The very highly significant differences between the PA of LD-derived and GWAS-derived marker we observed under the non-stressed E1 environment, suggest that the genetic architecture of the three phenotypic traits considered in our study deviates, to a certain degree, from the infinitesimal model, and that each trait is controlled by different sets of QTLs. Such deviations have been reported for the distribution of the effects of genes affecting quantitative traits in dairy cattle populations [62]. These authors performed a meta-analysis of large set of QTL mapping experiments and observed the maximum likelihood for a gamma distribution of QTLs effects, i.e. few QTL with large effects and many, but finite number, of QTLs with small effects, non-evenly distributed along the genome. The consequences of such deviation is that the relationship matrix G built with LD-based matrix does not provide an optimal description of the variance-covariance structure between individuals for the trait considered, leading to lower PA levels.

Gains in PA with S-GWAS-based marker selection were lower but more realistic in the context of actual breeding programs in which phenotypic data are not available for the individuals that are candidates for selection. Similar to GWAS-derived markers, the differences in gains in PA observed in S-GWAS-derived markers, between the population of 280 and 204 accessions, suggest that the size of the population plays a determining role in the informativeness of the trait-specific markers. In other words, GWAS performed with 224 accessions (the size of the training set for S-GWAS in the population of 280 accessions) is more informative than GWAS performed with 163 accessions, the size of the training set for S-GWAS in the population of 204 accessions. This is in agreement with the well-known positive relationship between the size of the population and the power of GWAS to accurately evaluate the effect of each marker [63].

### Predictive ability of multi-environment predictions

The PA of genomic predictions obtained with the multi-environment models were, on average, similar to their single environment counterparts under the CV1 cross-validation strategy, and significantly higher under the CV2 cross-validation strategy. Gains in the PA of the multi-environment models combined with CV2 were the highest in the two drought affected environments, E2 and E3. The two multi-environment models tested (GBLUP, RKHS) led to significantly different predictive abilities. Under GBLUP, the overall average gain in PA was 7% and ranged from – 4% for GY to 15% for PH. These gains were significantly lower than the 30% gain reported in [38] comparing multi-environment and single environment GBLUP models for GY in wheat. This was also the case for the gains in PA reported in [42] using rice data from two managed environments, continued flooding and alternate watering and drying. These authors reported gains of up to 29% for DTF compared with 22% in our case. The RKHS multi-environment model enabled gains in PA of up to 32%. These gains are much lower than those of up to 68% reported in [39] in wheat, similar to the ones reported by [42], and in accordance with [41], higher than the GBLUP multi-environment model. The differences in the amplitude of gains of PA observed in our study and the ones reported in [39, 41] are probably due to several factors, including the size of the population (599 wheat lines against 204 accessions in our case), the number of environment (large number of environments that were grouped in four target sets of environments versus three in our case), and the distinctive features of those environments. While in the case of wheat, data from a multi-local trial across a natural continuum of environments were grouped in four subsets, our data were produced in a managed-environment trial with clear-cut treatments to assess the effect of drought stress on rice development end yield. Overall, these results reinforce the conclusion drawn by [42] that, multi-environment genomic prediction models that account for G×E interactions as evaluated from multi-local trials, are also of interest for breeding for tolerance to abiotic stresses, including drought.

### Implications for breeding rice for drought tolerance

Improving resilience to drought during floral development and anthesis is an important target for major cereal crops including rice [1, 64]. It is now widely accepted that the ability to predict complex traits is better using whole-genome marker prediction than using a few markers that target a few quantitative trait loci [65, 66]. In the case of rice, several empirical studies embedded in ongoing breeding programs have confirmed the potential of GS in accelerating genetic gains [34, 36, 42]. Recently, a new generation of genomic prediction models was developed that further improve the PA of genomic prediction by explicitly modeling G×E interactions in the same way as in multilocal trials that all breeding program resort to [37-41]. The effectiveness of such models in the context of breeding for tolerance to abiotic stresses was first documented in rice [42] using data from managed-environment trials to evaluate the performances of a reference population and a progeny population under two water management systems: alternate watering and drying and conventional continuous flooding. These authors also reported gains in PA of up to 14% compared to direct selection based on phenotypic correlation between the two water management systems. Likewise, they showed that the share of non-phenotyped individual in the CV2 prediction strategy could be increased up to 40% with no significant negative effect on PA.

In the present study, we showed that (i) reasonably high PA of genomic prediction can be obtained with as few as 3,000 SNP markers, even in a population of limited LD extent, provided the markers are selected on the basis of LD. (ii) Trait-specific markers resulted in higher PA of genomic prediction than markers selected on the basis of neutral LD, especially when the size of the training population is large enough to allow the accurate estimation of the effect of the markers. (iii) Tolerance to drought stress was better predicted by multi-environment models (especially RKHS) that accounted for G×E interaction (E being non-stressed lowland and stressed lowland or upland, or both), than their single-environment model counterparts, when associated with the CV2 prediction strategy. (iv) Finally, yet importantly, even under the less favorable CV1 prediction strategy, multi-environment models achieved similar PA to their single environment counterparts.

The final finding suggests the feasibility of a genomic breeding scheme aiming at simultaneous improvement of yield potential and drought tolerance. It requires a training population carefully phenotyped under both favorable environmental conditions and managed drought, while in the first step of selection, the candidate population would be phenotyped only under favorable environments. The selected candidate would be phenotyped under managed drought to ascertain their GEBV and to update the multi-environment prediction model for the next breeding cycle [67-68]. The process can be implemented in the framework of the pedigree-breeding of progeny of biparental crosses between members of the reference population of the breeding program that is used as the training population [36; 42]. In the context of multi-environment model-based genomic predictions, the effectiveness of trait-specific markers merits investigations using a simulation approach. Nevertheless, breeders should consider including a limited share of trait specific markers (especially for the most important target traits) when genotyping their candidate populations.

## Figures and Tables

**Table 1:** Sources of phenotypic variation and derived summary statistics of days to flowering (DTF), grain yield (GY) and plant height (PH) under three drought environments, E1, E2 and E3.

Single environment analysis
| Trait | Drought environment | Mean | SD | Phenotypic variances |  |  |  | H<sup>2</sup> | SE | Phenotypic correlation |  |  |
| --- | --- | --- | --- | --- | --- | --- | --- | --- | --- | --- | --- | --- |
|  |  |  |  | Fixed effects | Random effects |  | Total |  |  | E1-E2 | E2-E3 | E1-E3 |
|  |  |  |  |  | Genotype | Residual |  |  |  |  |  |  |
| DTF | E1 | 86.7 | 9.53 | 0.38 | 70.27 | 23 | 93.64 | 0.75 | 0.05 | 0.71* |  |  |
|  | E2 | 85.84 | 38.3 | 0 | 211.17 | 67.41 | 278.58 | 0.76 | 0.03 |  | 0.58* |  |
|  | E3 | 90.19 | 32.44 | 0 | 348.52 | 54.55 | 403.07 | 0.84 | 0.01 |  |  | 0.68* |
| GY | E1 | 4.98 | 1.68 | 4.26 | 3,425.56 | 1,578.10 | 5,007.92 | 0.68 | 0.07 | -0.18* |  |  |
|  | E2 | 0.18 | 0.19 | 2.14 | 50.22 | 40.15 | 92.51 | 0.54 | 0.05 |  | 0.50* |  |
|  | E3 | 0.43 | 0.62 | 1.77 | 137.38 | 11.98 | 151.12 | 0.91 | 0.08 |  |  | -0.16* |
| PH | E1 | 106.56 | 21.2 | 0 | 469.94 | 21.71 | 491.65 | 0.96 | 0.01 | 0.64* |  |  |
|  | E2 | 68.57 | 12.38 | 0.84 | 199.19 | 83.17 | 283.2 | 0.71 | 0.03 |  | 0.74* |  |
|  | E3 | 73.62 | 13.03 | 0.84 | 194.98 | 45.36 | 241.18 | 0.81 | 0.02 |  |  | 0.59* |

Combined analysis of the three environments
|  | Phenotypic variance |  | ZValue |  | ProbZ |  |  |  |
| --- | --- | --- | --- | --- | --- | --- | --- | --- |
|  | Genotype | Residual<sup>(1)</sup> | Genotype | Residual | Genotype | Residual |  |  |
| DTF | 274.58 | 274.58 | 9.16 | 28.55 | <.0001 | <.0001 | 0.76 | 0.05 |
| GY | 208.38 | 208.38 | 7.06 | 28.55 | <.0001 | <.0001 | 0.71 | 0.06 |
| PH | 178.74 | 178.74 | 9.6 | 28.55 | <.0001 | <.0001 | 0.96 | 0.01 |

SD: Standard deviation; Fixed effects: Bloc(Rep); H<sup>2</sup> : Broad sense heritability for single environment analysis; SE: Standard error for H<sup>2</sup>. \*: p-value of correlation <0.01. In the combined; (1): The residual variance includes the genotype by environment interaction.

## Supplemental information

**S1 Table**: List and main characteristics of the 280 rice accessions used in this study

**S2 Table**: Size of incidence matrices obtained by combining three levels of minor allele frequency (MAF) and five levels of linkage disequilibrium (LD)

**S3 Table**: (A) Predictive ability (PA) of genomic prediction obtained in cross validation experiments within the population of 280 rice accessions using GBLUP and RKHS prediction methods and 15 incidences matrices combining 3 levels of minor allele frequency (MAF) and five levels of Linkage disequilibrium (LD). Mean predictive ability (PA) and standard deviation (STD) of 100 cross-validation replicates are presented. (B) Analysis of factors that influence the variation in predictive ability.

**S4 Table**: Analysis of effect of marker selection methods on predictive ability of genomic predictions, in the population of 280 accessions. (A): Comparison of Linkage disequilibrium (LD) and GWAS based marker selection. (B): Comparison of Linkage disequilibrium (LD) and S-GWAS based marker selection.

**S5 Table**: Analysis of effect of marker selection methods (LD-derived and GWAS-derived markers) on predictive ability of genomic predictions, in the population of 204 accessions.

**S6 Table**: Analysis of effect of marker selection methods (LD derived and S-GWAS-derived markers) on predictive ability of genomic predictions, in the population of 204 accessions.

**S7 Table**: Predictive ability (PA) obtained with single-environment and multi environment models under 2 cross validation strategies (CV1 and CV2) and two prediction method (GBLUP and RKHS) with the population of 204 rice accessions, evaluated under three environments (E1, E2 and E3).

**S1 fig.** Heat map for the incidence matrices with 215,250 and 28,091SNP markers revealing an uneven distribution of markers along the rice genome.

**S2 fig.** Patterns of decay in linkage disequilibrium in the population of 280 accessions genotyped with 28,091 SNP.The curve represents the average r^2^ according to pairwise distance between markers among the 12 chromosomes and the bars represent the associated standard deviation.

**S3 fig.** Unweighted neighbor-joining tree based on simple matching distances constructed from the genotype of 280 accessions, using 4,824 SNP markers. Green: accessions belonging to *aus* group. Other colors: different subgroups of *indica* as established in the 3K rice genome project [45].

## Availability of data and materials

The Phenotypic data analyzed for this study are included in the supplementary table 1

The genotypic datasets generated and analysed during the current study are available in hapmap format at the following address http://tropgenedb.cirad.fr/tropgene/JSP/interface.jsp?module=RICE, in the tab “STUDIES” then selecting for “Genotypes” in *Study type* and for “GS-Ruse_IRRI-Reference-Population.

## Authors’ contributions

NA and AK conceived the study. AB produced the phenotypic data and contributed to data analysis. NA, JB and TVC analyzed the data. NA wrote the manuscript.

### Acknowledgments

This work was funded by Agropolis Foundation (http://www.agropolis-fondation.fr/) and Cariplo Foundation (http://www.fondazionecariplo.it/), Grant no 1201-006. This work was supported by the CIRAD - UMR AGAP HPC Data Center of the South Green Bioinformatics platform (http://www.southgreen.fr/)

## Conflict of Interest

The authors declare that they have no conflict of interest.

